# Kinetics characterization of ASXL1/2-mediated allosteric regulation of BAP1 deubiquitinase

**DOI:** 10.1101/2020.02.24.962464

**Authors:** Hongzhuang Peng, Joel Cassel, Daniel S. McCracken, Jeremy W. Prokop, Paul R. Collop, Alexander Polo, Surbhi Joshi, Jacob P. Mandell, Kasirajan Ayyanathan, David Hinds, S. Bruce Malkowicz, J. William Harbour, Anne M. Bowcock, Joseph Salvino, Eileen J. Kennedy, Joseph R. Testa, Frank J. Rauscher

## Abstract

BAP1 is a ubiquitin hydrolase whose deubiquitinase activity is mediated by polycomb group-like protein ASXL2. Cancer-related mutations/deletions of *BAP1* lead to loss-of-function either by directly targeting the catalytic (UCH) or ULD domains of BAP1, the latter disrupts binding to ASXL2, an obligate partner for BAP1 enzymatic activity. However, the biochemical and biophysical properties of the domains involved in forming the enzymatically active complex are unknown. Here we investigate the molecular dynamics, kinetics and stoichiometry of these interactions. We demonstrate that the BAP1 and ASXL2 domain/proteins or protein complexes produced in either bacteria or baculovirus are structurally and functionally active. The interaction between BAP1 and ASXL2 is direct, specific, and stable to *in vitro* biochemical and biophysical manipulations as detected by isothermal titration calorimetry, GST association, and optical biosensor assays. Association of the ASXL2-AB box greatly stimulates BAP1 deubiquitinase activity. A stable ternary complex can be formed comprised of the BAP1-UCH, BAP1-ULD, and ASXL2-AB domains. Binding of the BAP1-ULD domain to the ASXL2-AB box is rapid, with fast association and slow dissociation rates. Stoichiometric analysis revealed that one molecule of the ULD domain directly interacts with one molecule of the AB Box. Real-time kinetics analysis of ULD/AB protein complex to the UCH domain of BAP1, based on SPR, indicated that formation of the ULD/AB complex with the UCH domain is a single-step event with fast association and slow dissociation rates. These structural and dynamic parameters implicate the possibility for future small-molecule approaches to reactivate latent wild-type UCH activity in BAP-mutant malignancies.

## Introduction

BAP1 was discovered as an ubiquitin hydrolase that associates with the RING finger domain of BRCA1 and enhances BRCA1-mediated inhibition of breast cancer cell growth (1). The N-terminus of BAP1 consists of a UCH domain (ubiquitin C-terminal hydrolase) that cleaves ubiquitin from ubiquitin-conjugated small substrates. BAP1 contains two protein-binding motifs for BARD1 and BRCA1, which form a tumor suppressor heterodimeric complex (2), and a binding site for HCF1, which interacts with a histone-modifying complex during cell division (3). The C-terminus of BAP1 contains two nuclear localization signals and ULD (UCH37-like domain). The ULD domain interacts with ASXL family members to form the polycomb group (PcG)-repressive deubiquitinase complex involved in stem cell pluripotency and other developmental processes (4, 5).

Homology of the BAP1-UCH and other UCH-like proteins implies a role for either ubiquitin-mediated, proteasome-dependent degradation or other ubiquitin-mediated regulatory pathways in BRCA1 function, in cellular growth, differentiation, and homeostatic processes (1, 6, 7). BAP1 exhibits tumor suppressor activity in cancer cells (1, 2) and *in vivo* (8). Moreover, not only were somatic mutations/deletions of *BAP1* found in metastasizing uveal melanomas, malignant mesothelioma, and other cancers (9-11), but also germline mutations of *BAP1* were found in families with a high incidence of mesothelioma, uveal melanoma, benign and malignant cutaneous melanocytic tumors, basal cell carcinoma, meningioma, and renal carcinoma (11-15). Cancer-related mutations/deletions of *BAP1* often result in loss-of-function by causing premature protein termination, protein instability and/or loss of UCH catalytic activity. Other mutations of *BAP1* lead to loss-of-function by targeting the ULD domain, thereby disrupting binding to ASXL2 (16) an obligate partner for BAP1 enzymatic activity.

BAP1 functions as part of a large polycomb-like complex throughout vertebrate and invertebrate biology through the ASXL1/2 family members (5). The *Drosophila* PcG Calypso protein is homologous to BAP1. Calypso interacts with PcG protein *Asx*, and this Polycomb repressive deubiquitinase (PR-DUB) complex binds to PcG target genes. The human homologs of *Asx* are *ASXL1-3* (16). The N-terminus of ASXL contains the highly conserved *Asx* homology domain (ASXH), which is required for Calypso/BAP1 protein binding. Similar to *Drosophila Asx*, human ASXL1/2-BAP1 complexes deubiquitinate histone H2A. Mutations of *ASXL1/2/3* genes leading to protein truncations have been found associated with human cancers and other diseases (17-20). One example is loss-of-function mutations in *ASXL1*, which encodes an epigenetic modifier that plays a role in polycomb repressive complex (PRC2)-mediated transcriptional repression in hematopoietic cells. Such loss-of-function mutations in myeloid malignancies result in loss of PRC2-mediated gene repression of leukemogenic target genes (17). The crystal structure of *Drosophila* PR-DUB, has revealed that the deubiquinase Calypso and its activating partner ASX form a 2:2 complex. This bidentate Calypso ASX complex is generated by dimerization of two activated Calypso proteins through their coiled-coil regions. Disrupting the Calypso dimer interface does not affect inherent catalytic activity, but inhibits removal of H2AK119Ub as a consequence of impaired recruitment to nucleosomes (21).

In a early previous study, we found that the familial and somatic *BAP1* mutations inactivate ASXL1/2-mediated allosteric regulation of BAP1 deubiquitinase by targeting multiple independent domains (16). The AB Box of ASXL2 mediates the binding of ULD and UCH domains of BAP1 to form a tripartite complex, which subsequently stabilizes the UCH structure, thereby increasing the catalytic activity of BAP1-UCH. The tumor-derived discrete in-frame deletions and insertions outside of the BAP1 catalytic domain (UCH) disrupt the BAP1-ASXL2 interaction, leading to tumor-related loss of BAP1 catalytic activity. In this study, we define the biochemical and biophysical properties of the domain-domain interactions of this complex. Importantly, these new studies elucidate the molecular dynamics of the interaction, measure the kinetic and stoichiometric impact of mutations on proteins binding and on the enzymatic activity of BAP1, and provide novel insights about the structural and dynamic parameters of the BAP1-ASXL2 interaction into single cell datasets that inform future small-molecule approaches to reactivate latent wild-type UCH activity in BAP-mutant malignancies.

## Materials and methods

### Plasmids

The pFastBacTHa-BAP1-FL-WT, -BAP1-UCH-WT and -UCH-C91S mutant plasmids, pGEX-2TK-BAP1-UCH-WT (1-250 aa), pGEX-4T-1-BAP1-ULD, pQE30-BAP1-ULD (601-729aa) and pQE30-ASXL2-AB (261-381aa) plasmids were previously described (16). The pETDuet-1-His-BAP1-ULD+ASXL2-AB plasmid was constructed through PCR-based cloning and was sequenced to confirm its authenticity.

### Proteins expression and purification

The baculovirus (Bv) Bv-His-BAP1-FL-WT, Bv-His-BAP-UCH-WT and Bv-UCH-C91S mutant proteins were expressed in Bv-infected Sf9 cells and purified as previously described (16). The GST- and His-tagged BAP1 and ASXL2 proteins were expressed in *E. coli* BL21 (DE3) (Stratagene) and SG13009 (S9) (Qiagen), respectively. The pETDuet-1-His-BAP1-ULD+ASXL2-AB protein complex was expressed in Rosetta 2 (DE3) pLysS (Millipore). The bacteria bearing the desired plasmids were propagated with aeration at 37°C in 1L of 2YT to an A_600_ absorbance of approximately 0.6. IPTG was added to 1 mM, and growth was continued at 20°C overnight. The cells were harvested by centrifugation.

GST-fusion proteins were purified as described previously (22). The bacterial His-tagged proteins were purified under denaturing conditions (Qiagen) and refolded by dialysis as described previously (22). The recombinant human BAP1-FL-WT protein was purchased from Boston Biochem (E-345-050). The Duet-His-ULD/AB protein complex was purified under native purification conditions using Cobalt beads (Talon), followed by dialysis and concentration to desired concentration.

### GST association assays

GST association assays were performed as described previously (23) using BB200 buffer (200 mM NaCl, 20 mM Tris, pH 7.5, 0.2 mM EDTA, 10% Glycerol, 1 mM PMSF and 0.2% NP40) and BB500 (containing the same components as BB200 except that the concentration of NaCl was 500 mM).

### Dynamic light scattering (DLS)

DLS was measured using DynaPro Titan (Wyatt Technology). Purified His-BAP1-ULD, His-AB and His-ULD/AB complex were in buffer containing 50 mM potassium phosphate, pH 7.5, 200 mM potassium chloride and 1mM TCEP. His-ULD was measured at 574 µM concentration. His-AB was measured at 77 µM concentration. The His-ULD/AB protein complex was measured at 70 µM□□□□centration. Samples were spun in a tabletop microcentrifuge at 13,000 rpm for 10 minutes prior to measurements, and measurements were done at 10°C.

### Isothermal titration calorimetry (ITC)

ITC was performed using Microcal ITC 200 (Microcal/Malvern Instruments). His-ULD and His-AB proteins were dialyzed in 50 mM potassium phosphate, pH 7.5, 200 mM potassium chloride, and 1 mM TCEP. His-AB was placed in the sample cell at concentration 77 µM. His-ULD was titrated into the sample cell at a concentration of 574 µM. Two references were used. The first reference was titration of the buffer into His-AB protein. The second reference was titration of His-ULD protein into the buffer. Both reference values were subtracted from the experimental data. ITC calculations and fitting was performed with Origin 7 software, using autofit, 200 iterations. Based on the results, the stoichiometry and binding kinetics of the proteins were determined. The direct measurements of binding affinity (K_a_), enthalpy changes (ΔH) and binding stoichiometry (n) were used to determine the Gibbs free energy changes (ΔG) and entropy changes (ΔS) using *ΔG=-RTlnK*_*a*_*= ΔH-TΔS* (R = gas constant; T = absolute temperature). Dissociation constant (K_d_) is 1/K_a_. Experiments were performed in duplicate. No uncertainty ranges are given due to the low number of technical replicates.

### Circular Dichroism (CD)

The CD spectra (190-260 nm) were measured on a Jasco J-715 spectropolarimeter (Japan Spectroscopic Co.) at 25°C. The CD spectra were recorded using 0.1 cm path length quartz cuvette with the following measurement parameters: 190-260 nm; step resolution: 1 nm; speed: 20 nm/min; accumulations: 4; bandwidth: 1 nm. All measurements were performed in the following buffer: 50 mM potassium-phosphate, pH 7.5, 300 mM KCl, 10% glycerol, 1 mM DTT and 1 mM PMSF. The data were processed using the Jasco Spectra Manager Suite.

### Ub-AMC assay

The activity of BAP1 or BAP1-UCH proteins was determined by cleavage of ubiquitin-7-amido-4-methylcoumarin (Ub-AMC). Assays contained various concentrations of enzyme and substrate with and without His-AB or the His-ULD/AB complex as indicated in the figures in a reaction volume of 15 uL of 25 mM HEPES, pH 7.4, 150 mM NaCl, 5 mM DTT, 0.005% Tween20 in low-volume 384-well plates at room temperature. Fluorescence of free AMC at excitation and emission wavelengths of 355 nm and 460 nm, respectively, was measured at 2 min intervals for 20 min in an Envision microplate reader. Background fluorescence in the absence of enzyme was subtracted from the data points, and the linear portion of the curve was fit to a straight line to determine velocity.

### Kinetic analysis: surface plasmon resonance (SPR)

Interactions between the ASXL-AB and BAP1-ULD domains were studied by SPR using a Biacore T200 instrument. GST-antibody (Abcam ab9085) was coupled to all flow cells of a CM5 sensor chip using standard amine coupling procedures in a HEPES-buffered saline running buffer. After coupling of the GST antibody, the running buffer was changed to 25 mM HEPES, pH 7.4, 150 mM NaCl, 5 mM DTT and 0.05% Tween20. GST-ULD was immobilized onto the chip surface at a ligand density of 400 RU, followed by a 120-s stabilization period. A single concentration His-AB was then injected over both the reference cell, with GST antibody alone, and the flow cell covered with GST-ULD at 30 uL/min. The binding reaction was monitored for 240 s followed by a 300-s dissociation time. Specific binding was determined by subtracting the refractive index change in the reference cell from the flow cell containing GST-ULD. After each concentration of His-AB, the GST-ULD was stripped from the surface using a 60-s injection of 20 mM glycine, pH 2.0 at 30 uL/min, followed by another 120-s stabilization period. Fresh GST-ULD was then immobilized as above. Experiments were done in triplicate.

Interactions between the His-ULD/AB complex and full-length BAP1 or the BAP1-UCH domain were also studied using the Biacore T200 instrument. Full-length His-BAP1, GST-UCH, or GST alone was directly immobilized to a CM5 sensor chip at a density of ∼3000 RU using standard amine coupling procedures. The running buffer for the binding studies was 25 mM HEPES, pH 7.4, 250 mM NaCl, 5 mM DTT and 0.05% Tween20. The higher NaCl concentration was required to reduce nonspecific binding to the reference cell in the absence of protein. Various concentrations of His-ULD/AB complex were injected over the flow cells at 30 uL/min and the binding reaction monitored for 90 s followed by a 240-s dissociation time. Specific binding was determined by subtracting the refractive index change in the reference cell from the readings of the other three flow cells. After the 240-s dissociation time, most of the His-ULD/AB complex was completely dissociated. However, 1 M NaCl at 30 uL/min was injected for 60 s over the flow cells to clear any remaining bound protein. Experiments were done in triplicate.

### Sequence and structure analysis

Open reading frame sequences for BAP1, UCHL1, UCHL3, UCHL5, ASXL1, ASXL2, and ASXL3 were obtained from NCBI for vertebrate species. Separately, the UCH or ASX sequences were aligned and codon selection scored using our previously published metrics (PMID: 28204942). COSMIC variants (PMID: 25355519) for BAP1 were extracted on June 20, 2018. Secondary structure predictions for proteins were performed using http://cib.cf.ocha.ac.jp/bitool/MIX/, a combination of Chou-Fasman, GOR, and Neural Network predictions. Conservation was highlighted onto the human protein model generated from PDB 6cga.

## Results

### Bap1 and Asxl protein coexpression in single cell RNAseq datasets

To build a cellular model of *Bap1* and *Asxl1-3* coexpression, we used the 53,760 cell dataset of 20 tissues from mouse (PMID: 30283141). *Bap1* expression was found to vary in the average counts per cell and the number of cells expressing the gene, with tissues such as thymus showing the highest *Bap1* levels (Suppl. Fig. S1A). Co-segregating gene expression in those cells expressing *Bap1* versus those that do not for the thymus revealed 122/289 genes that positively correlated to be involved in cell cycling (p-value, 3.5e-61) and several that were connected to BAP1 interaction pathways (Suppl. Fig. S1B). Interestingly, cancer-related genes such as *Fos* were negatively correlated with *Bap1*. Among the 20 tissues, the majority of *Bap1*-expressing cells had none of the *Asxl1-3* genes expressed (60.7%), with 21.1% of cells repressing *Asxl2*, 11.3% of cells with *Asxl1*, 5.9% of cells expressing both Asxl1 and Asxl2, and 0.9% with AsxL3 (Suppl. Fig. S1C), suggesting ASXL2 kinetic interactions are of the highest priority for ASXL proteins.

The breakdown of the 20 tissues showed a varying percentage of *Bap1* positive cells to have *Asxl1* or *Asxl2* expression, with tissues such as pancreas having the greatest *Asxl2* bias and those such as muscle having an *Asxl1* bias (Suppl. Fig. S1D). Correlation analysis of the single cells for each tissue revealed that liver and pancreas have higher correlations between *Asxl2* and *Bap1* expression levels (Suppl. Fig. S1E), with genes correlating to those *Bap1* and *Asxl2* positive-expressing cells having significantly enriched protein-protein interactions (PPI) and lipid metabolic process gene ontology (GO) for positively correlated genes and regulation of cell motility in negatively correlated genes (Suppl. Fig. S1F).

### Analysis of conserved and selected BAP1 and ASXL1-3 contact sites

The domain structure of BAP1 is unique from other UCH proteins (Fig. 1A). The N-terminus of BAP1 has similarity to other mammalian UCHs (UCH-L1, UCH-L3, and UCH-L5); however, BAP1 also has several additional conserved motifs and domains throughout the remainder of the protein including the ULD found only in UCHL5. Alignments of the UCH domain of the four proteins and the ULD of BAP1 and UCH-L5 identify many amino acids conserved throughout, especially at sites with cancer (COSMIC) mutations within the UCH (Fig. 1B-C). Using the structure of *Drosophila* Calypso UCH/ULD interaction with Asx (PDB 6HGC) converted into human BAP1 UCH/ULD and ASXL2 merged with our previous models of interaction with H2A and Ubiquitin (16), we can pinpoint the human contact maps of the ULD with ASXL2 with high confidence (Fig. 1D, E). The BAP1 ULD contact amino acids have 12/22 amino acids fixed throughout the evolution of both BAP1 and UCH-L5, yet 7/22 amino acids are unique to BAP1, suggesting a lower kinetics of interaction between ASXL2 and UCH-L5 than with BAP1. The conservation of ASXL1-3 identifies a shared highly conserved ASXH domain critical for ULD interaction, with an additional conserved PHD-type domain being poorly defined (Fig. 1F). Of the BAP1 contact amino acids within ASXL2, 20/29 sites are conserved between *Drosophila* ASX and ASXL1-3 (Fig. 1G). A total of 23/29 BAP1 contact sites are conserved throughout ASXL1-3, suggesting that contact between ASXL1-3 with BAP1 are maintained throughout all three proteins.

**Figure 1.**
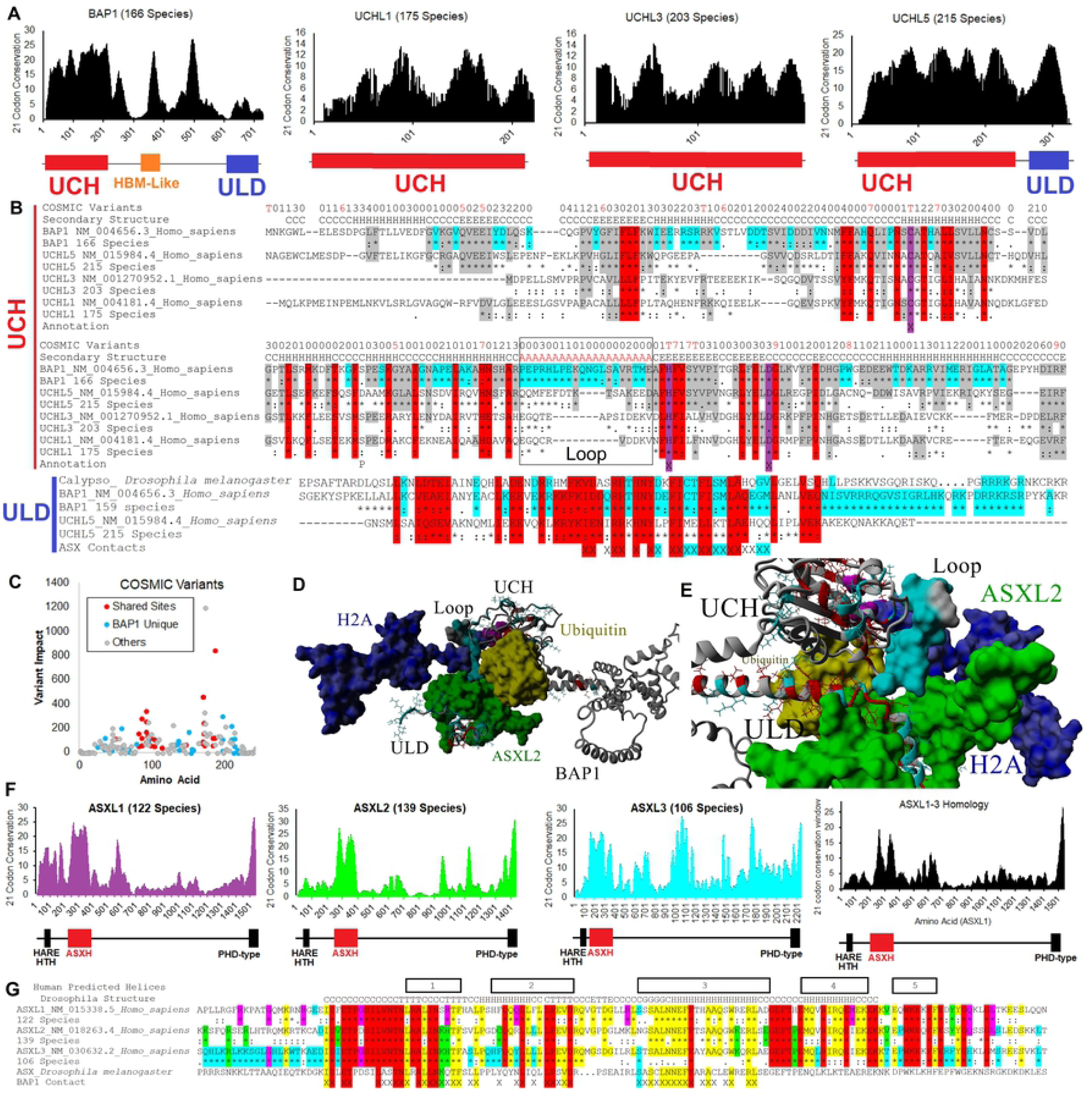
BAP1 structure and evolution. **A)** Open reading frame sequences were aligned for each UCH protein followed by assessment of amino acid conservation and codon selection. Number of each species sequences used is listed next to each name. The scores for each site were placed on a 21-codon sliding window, adding scores for 10 up- and down-stream of any site. Annotated domains are shown below each. **B)** Sequence alignments of UCH domain (red line) or ULD (blue line) of BAP1, UCHL5, UCHL3, and UCHL1 showing the human sequence of each with the consensus alignment information below each (* = conserved in all species for each gene: = functionally conserved for each gene; = weakly conserved in each gene). Shown on the top is the number of COSMIC variants observed at each site (T = values ≥10) and below that is the secondary structure annotated based on protein modeling of the UCH. The X in annotation marks amino acids in the enzyme active site. Amino acids highlighted in red are conserved in all sequences, those in gray are conserved in at least two different proteins, and those in cyan are conserved and unique to BAP1. Sequence alignment of the ULD of BAP1 and UCHL5 includes the Drosophila Calypso sequence and the ASX contact amino acids marked with X. **C)** COSMIC variants of the UCH annotated for variant impact and based on conservation with other UCH proteins with coloring based on panel B. **D-E)** Model of the BAP1-UCH domain with colors shown from the previous alignments with additional bound H2A (blue) with ubiquitin (yellow), and the ULD (conservation based on alignment in panel Fig. 1B) revealing additional BAP1 uniquely conserved amino acids for the stabilization by ASXL2 depicted in green near the UCH loop (cyan). The entire complex is shown in panel C and a zoom in view of ASXL2, ULD, and UCH interactions in panel D. **F)** Open reading frame sequences aligned for ASX1-3 proteins followed by assessment of amino acid conservation and codon selection of all three combined (black). **G)** Sequence alignments of the AB boxes of ASXL1, ASXL2, and ASXL3, and *Drosophila* Asx. Amino acids in red are conserved in all three proteins, those in yellow shared in all three human sequences, those in magenta conserved at least in ASXL1, those in green at least in ASXL2, and those in cyan at least in ASXL3. Contact amino acids with BAP1 are marked with X.

Of note, BAP1-UCH contains a larger loop than the other UCH proteins, with a high conservation of both these loop amino acids and of multiple amino acids structurally near this loop (Fig. 1B, D, E), suggesting that larger substrates would be accessible to the catalytic cleavage site only for BAP1 and not other UCH domains, yielding a BAP1-specific recruitment of proteins/domains such as Asx and ULD for enzyme regulation. ASXL functions as a molecular scaffold to recruit BAP1 to transcription factors, which specifically bind to its target genes. Then, BAP1 ubiquitin hydrolase specifically removes the ubiquitin from histones of chromatin to regulate these target genes. ASXL not only functions as a molecular scaffold for BAP1 but also greatly stimulates its activity. When mutations/deletions occur in *BAP1*, either they cause enzymatic loss-of-function of BAP1, or abolish BAP1’s association with ASXL. Loss of binding to ASXL would dramatically decrease BAP1 deubiquitination activity, because of an inability to bring ASXL to BAP1’s catalytic site. On the other hand, products of *ASXL* gene mutations that lose association with BAP1 also lead to BAP1 loss of function. The structure of the BAP/ASXL2 tripartite complex has not been determined; however, the crystal structure of the *Drosophila* Calypso and its activating partner Asx was recently determined (21). The stoichiometry of BAP and ASXL1-3 interaction and the kinetics remained unknown. Therefore, we initiated biochemical and biophysical analyses of the BAP1-UCH, BAP1-ULD, ASXL2-AB domains and protein complex.

### Purification of recombinant proteins and protein complex

For single protein expression, His- or GST-tagged full-length BAP1, BAP1-UCH and BAP1-ULD domains were expressed in bacteria (Bac-) or baculovirus (Bv-), respectively (Fig. 2A). The reasons that we expressed the proteins in baculovirus were in case post-translational modifications are needed for the protein functions and/or that other cellular factors are involved in the protein functions. All the baculovirus-expressed proteins and domains were soluble using Ni^2+^-NTA chromatography under native purification conditions (Fig. 2B) and the proteins were functional (see below). The bacterial-expressed GST-BAP1-UCH and GST-BAP1-ULD were soluble using GST-chromatography under native purification conditions (Fig. 2B) and the proteins were functional (see below). The bacterial-expressed His-BAP1-ULD and His-ASXL2-AB proteins were purified under denaturing conditions, followed by a re-naturation protocol that yielded soluble, highly active proteins (Fig. 2B). However, the yield of re-folded proteins was not sufficient for structural studies. We thus used the pETDuet co-expression system to co-express His-ULD and AB, or His-AB and ULD protein complexes in *E. coli* [Rosetta 2 (DE3) pLysS]. The His-ULD/AB protein complex was successfully co-expressed and then purified using cobalt beads (Talon) under native purification conditions. The protein complex was highly soluble and functional (Fig. 2B).

**Figure 2.**
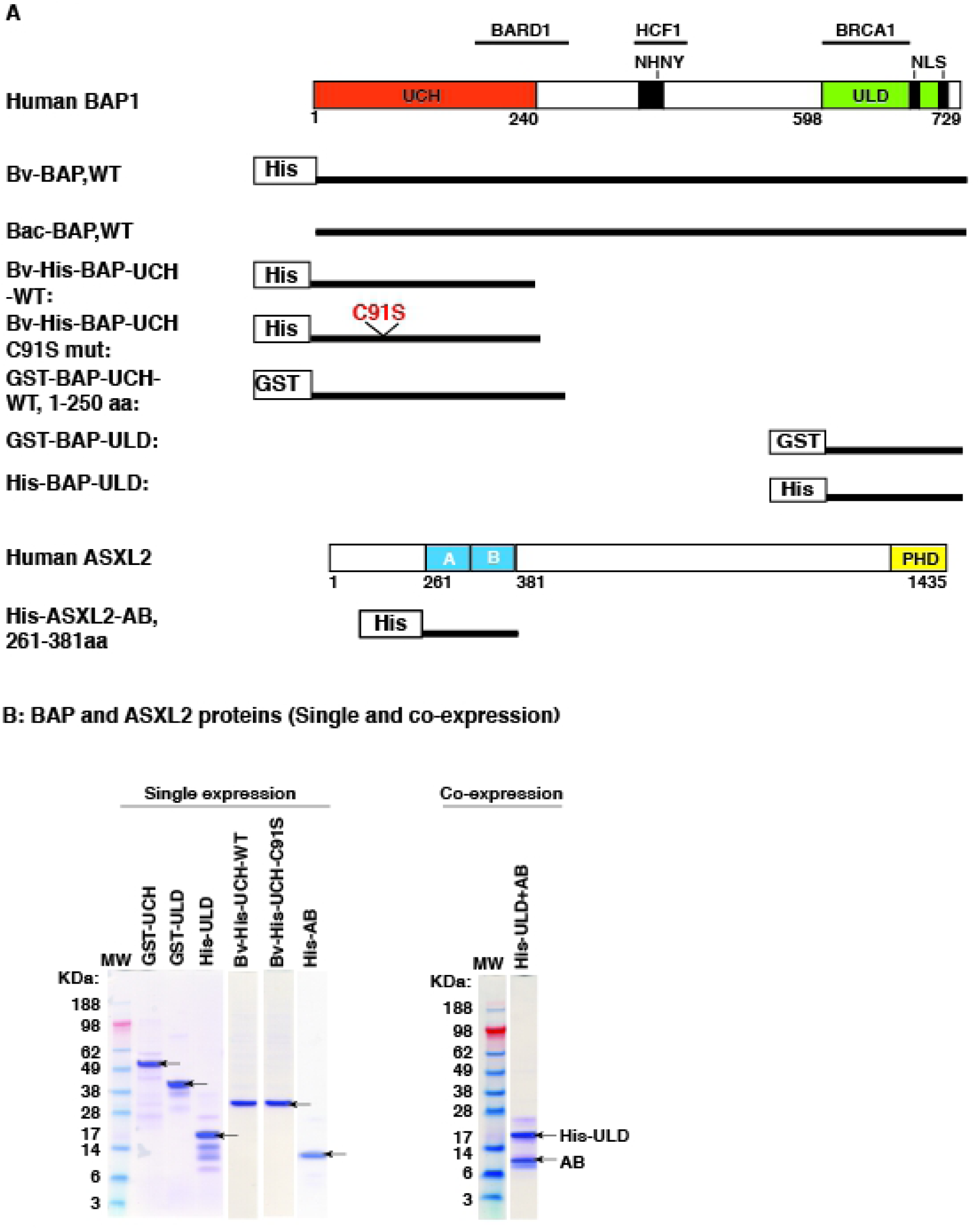
Domain architecture of human BAP1 and ASXL2 and the proteins/domains used in this study. **A)** Human BAP1 depicting ubiquitin C-terminal hydrolase domain (UCH; aa 1-240), BARD1 and BRCA1 binding domains, NHNY consensus sequence for interaction with HCF1, UCH37-like domain (ULD: aa 598-729), and nuclear localization signals (NLS). Domain structure of human ASXL2 contains highly conserved AB box and PHD domain. **B)** BAP1 and ASXL2 proteins produced in bacteria and baculovirus either singly or by co-expression. The proteins or protein complex were purified using either Ni-NTA, cobalt beads (Talon) or GST-resin. The purified proteins and protein complex were analyzed by NuPAGE and visualized by Coomassie stainining.

### Biophysical and biochemical characterization of BAP1-UCH, BAP1-ULD, ASXL2-AB, and the UCH/ULD/AB complex

To evaluate the behavior of singly-expressed proteins and co-expressed protein complex, DLS was used to examine the mono-dispersion of His-ULD, His-AB and His-ULD/AB complex. We first tested a full spectrum of buffer conditions for optimizing the solubility and stability of individual proteins and the protein complex. Under the optimal buffer condition found (50 mM potassium phosphate, pH 7.5, 200 mM potassium chloride and 1 mM TCEP), His-AB and His-ULD were mono-dispersed 87% and 88%, respectively. Each scan shows a larger species as well, which is assumed to be protein aggregation (Fig. 3A).

**Figure 3.**
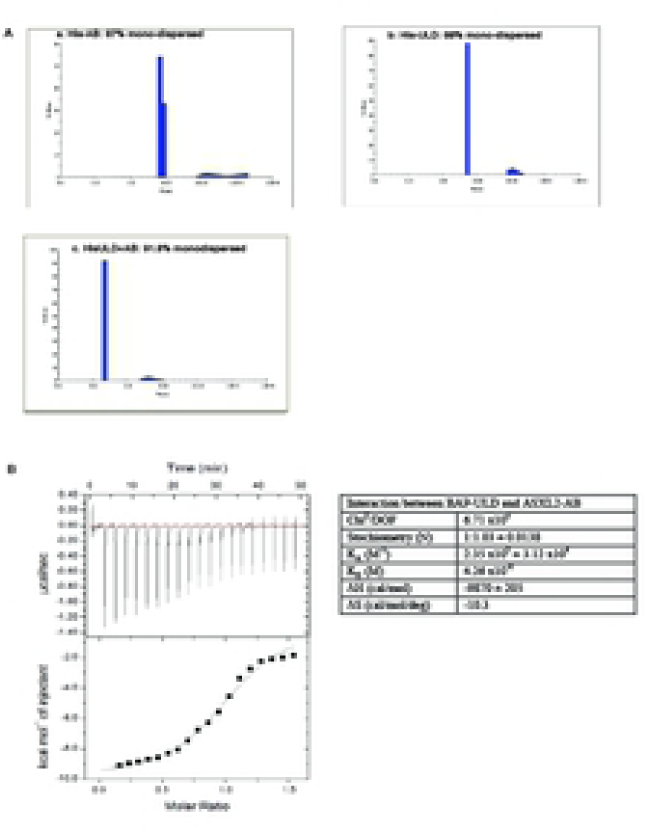
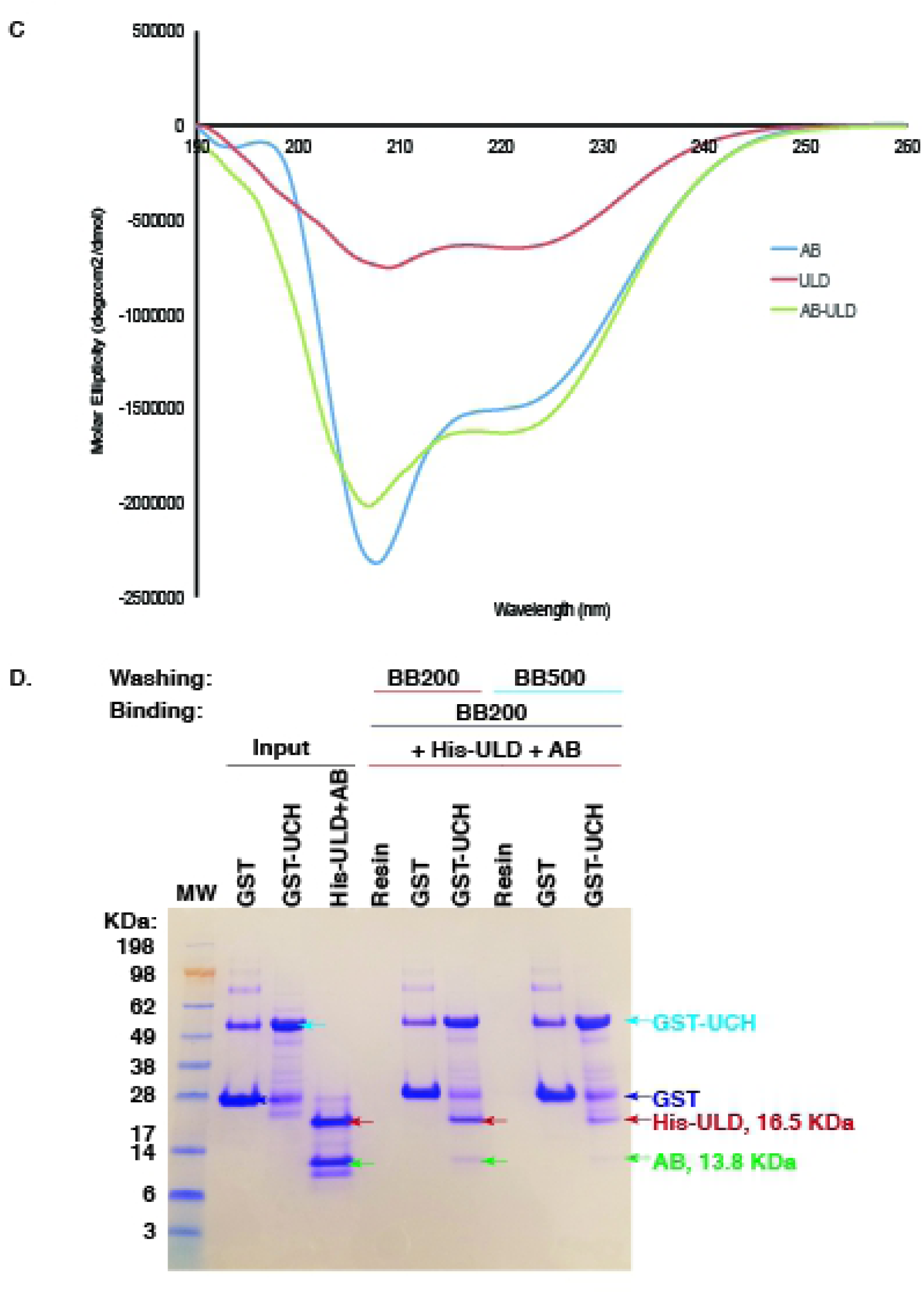
Biochemical and biophysical analyses of purified proteins and protein complex from BAP1 and ASXL2. **A)** Dynamic light scattering (DLS) was used to examine the mono-dispersion of His-ULD, His-AB and His-ULD/AB complex. Under the optimal buffer condition, His-ULD and His-AB proteins showed 88% and 87% mono-dispersion, while His-ULD/AB protein complex exhibited a higher degree (91.8%) of mono-dispersion, as directly measured by DLS. **B)** Isothermal titration calorimetry (ITC) was used to determine the thermodynamics and kinetics of domain-domain interactions between His-ULD and His-AB and their stoichiometry. 574 µM His-ULD protein was titrated into 77µM His-AB protein in terms of molar ratio. ITC calculations derived from the direct measurements and curve fitting were done with Origin 7 software. The binding affinity with dissociation constant of the protein-protein interaction and the stoichiometry of protein complex were determined. **C)** Circular dichroism was performed to determine the secondary structure of purified His-ULD and His-AB proteins as well as the His-ULD/AB protein complex. Data were processed using the Jasco Spectra Manager Suite. **D)** Binding of co-purified His-ULD/AB and the UCH domain of BAP1, as demonstrated by GST-UCH pull down with recombinant His-ULD/AB complex.

When this ULD-AB complex forms together, the mono-dispersion is measured at 91.8%. This indicates a similar, or perhaps slightly higher stability of the complex than the isolated proteins. In addition, we see a shift in the scan to a smaller size complex when these protein domains are bound together. This is contrary to what would typically be expected as proteins bind together. Based upon this result, it appears that the complex is more tightly packed spatially than the individual proteins. This result is consistent with the CD data presented, which show additional secondary structure formation attained during binding. In addition, these data were utilized for further ITC experiments (Fig. 3B) in calculating concentrations used, because it is assumed only the mono-dispersed species is capable of interacting properly with the other complex members.

From our previous studies (16), we learned that the BAP1-ULD domain interacts directly with the ASXL2-AB box. However, the binding kinetics and stoichiometry of interaction of the ULD domain and the AB box remained unknown. Using ITC, we have now determined the thermodynamics, kinetics, and stoichiometry of this domain-domain interaction. Highly purified His-ULD and His-AB proteins were critically equilibrated in the same buffer (50 mM potassium phosphate, pH 7.5, 200 mM potassium chloride and 1 mM TCEP). The His-AB was placed in the ITC cell with 77 µM protein concentration while the titrated protein His-ULD was at 574 µM protein concentration. We also set the references for each protein (see Materials & Methods) for subtraction from the experimental data. The data show that K_d_ for interaction of His-ULD and His-AB is approximately 4.26 µM (3.73 µM-4.85 µM). The stoichiometry of His-ULD to His-AB is 1:1 molar ratio (Fig. 3B). We also observed that the thermodynamics of the interaction has a ΔH of −9.87 kcal/mol and ΔS of -10.3 cal/mol/deg, indicating an exothermic interaction. These data are consistent with our previous studies that used computer modeling technology to predict the molecular model of BAP1-ULD interacting with ASXL2-AB (16). The interaction for both ULD and AB has a modest binding affinity dissociation constant. This result is consistent with expectations of formation of a protein-protein complex in a reversible manner.

From our computer molecular modeling studies, BAP1-ULD is predicted to form a few long helices, while ASXL-AB box is predicted to form five helices (16). We performed CD to determine the secondary structure of the purified recombinant protein His-ULD, His-AB and His-ULD/AB complex. The CD spectra of the domains and complex demonstrating that each adopts a partially helical conformation and has a high degree of secondary structure (Fig. 3C), While His-AB appears to be partly unstructured as demonstrated by a broad minima at 208 nm, this minima is lessened in the His-ULD/AB complex. The complex also has increased alpha-helical content relative to the two monomer proteins as indicated by an increased minima at 222 nm.

### Direct interaction between BAP1-UCH, -ULD domains and ASXL2-AB domain

Using computer molecular modeling of UCHL5 structures, we predicted that the BAP1-ULD domain folds back to the BAP1-UCH catalytic domain and that the ASXL2-AB box stabilizes the UCH catalytic loop via a unique BAP1 mechanism not seen in other UCH proteins, allowing for ubiquitin to fit into the active site (Fig. 1D,E). The GST-UCH directly interacted with the ULD domain but did not directly interact with the AB box, while the ULD domain recruited the AB box so that they form a stable complex (16). Now, we have co-expressed and co-purified the His-ULD/AB domain complex using the pETDuet system, which allowed us to obtain well-folded protein complex (Fig. 2B). To test this highly purified protein complex, a GST association assay was performed. GST or GST-UCH was pre-coated on the GST resin, followed by incubation with His-ULD/AB complex. After washing with BB200 or BB500 buffer, the GST resin with protein complex was extracted, analyzed by SDS-PAGE, followed by Coomassie staining. The result showed that the His-ULD/AB complex was pulled down by GST-UCH but not by GST (Fig. 3D). The UCH/ULD/AB protein complex was indeed formed.

### Stimulation of BAP1 deubiquitinase activity by ASXL2-AB and ULD/AB complexes

In order to measure BAP1 deubiquitinase activity, we used the fluorogenic substrate Ubiquitin-AMC (Ub-AMC). The activity of the UCH domain of BAP1 was 5-fold greater than the full-length BAP1, with specific activity values of 358 ± 6.6 pmol AMC/min/pmol E and 73 ± 2.4 pmol AMC/min/pmol E, respectively (Fig. 4). For both full-length BAP1 and the UCH domain, a point mutation of the cysteine residue at position 91 completely abolished enzyme activity (Fig. 4), consistent with previous observations (16).

**Figure 4.**
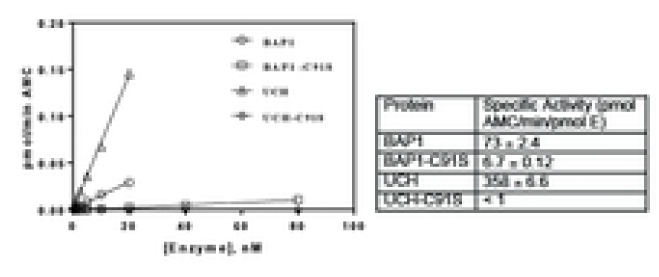
Cleavage of Ubiquitin-AMC mediated by full-length wild-type BAP1, full-length C91S BAP1 mutant, wild-type UCH domain of BAP1, or mutant C91S UCH domain. Enzymes were expressed in baculovirus with an N-terminal His-tag and purified using standard procedures. A range of concentrations for each enzyme was incubated with 100 nM Ubiquitin-AMC in 20 µL of 25 mM HEPES pH 7.4, 150 mM NaCl, 5 mM DTT and 0.005% Tween20 in 384-well plates. Fluorescence of free AMC was excited at 355 nm and emissions were measured at 460 nm at 2 min intervals. The resulting progress curves were fit to a straight line, and the velocities were plotted against enzyme concentration to obtain specific activities. Data points are means of duplicate determinations from a single experiment, which was repeated twice.

The ASXL-AB box stimulates BAP1 deubiquitinase activity in the Ub-AMC assay (16). In this study, we further characterized this effect by testing increasing concentrations of ASXL2-AB in the presence of a substrate titration of Ub-AMC. ASXL2-AB dose-dependently increased the maximal velocity of BAP1 cleavage of Ub-AMC by 2.5-fold (Fig. 5A). The K_m_ values for Ub-AMC in the presence of increasing concentrations of ASXL2-AB ranged from 4-9 mM and did not correlate with ASXL2-AB concentration, suggesting that the ASXL2-AB box stimulates BAP1 enzyme activity by increasing its V_max_, rather than the K_m_ for Ub-AMC. In addition, from these data we were able to obtain a functional potency for ASXL2-AB stimulation of BAP1 enzyme activity by plotting the V_max_ values for BAP1 enzyme activity against the concentration of ASXL2-AB box (Fig. 5B). These data fit well to a typical one-site dose response curve with a Hill slope of 1.0 and an EC_50_ of 0.96 nM (95%CI: 0.42-2.4 nM) (Fig. 5B).

**Figure 5.**
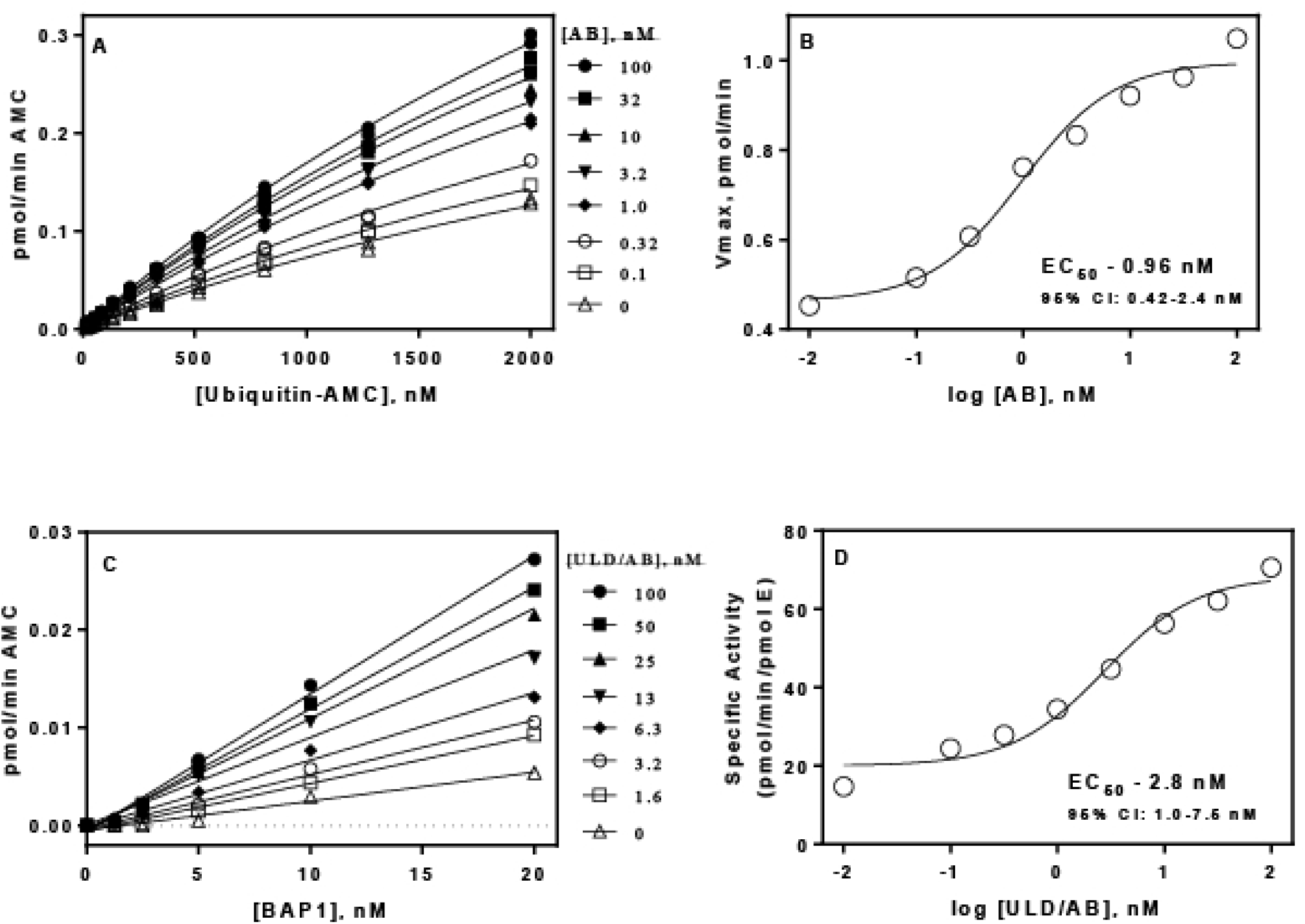
Effects of the AB box of ASXL2 and the ULD/AB complex of BAP1 mediated cleavage of Ubiquitin-AMC. **A)** Ubiquitin-AMC substrate titrations were incubated with full-length BAP1 (3 nM) in the presence of increasing concentrations of AB in assay buffer as described in Materials and Methods. The resulting progress curves were fit to a straight line and the velocities plotted against Ubiquitin-AMC concentration and the data fit to the Michaelis-Menton equation. **B)** Potency of AB-mediated stimulation of maximal velocity of BAP1. Each Vmax value from panel **A** was plotted against AB concentration, and the data fit to one-site dose response equation as described in Materials and Methods. **C)** Full-length BAP1 was titrated in the presence of increasing concentrations of ULD/AB complex and 100 nM Ubiquitin-AMC in assay buffer as described in Materials and Methods. The resulting progress curves were fit to a straight line, and the velocities were plotted against enzyme concentration to obtain specific activity. **D)** Potency of ULD/AB complex on specific activity of BAP1. Slopes from panel C were plotted against ULD/AB concentration and the data fit to one-site dose response equation as described in Materials and Methods. Data points are means of duplicate determinations from a single experiment, which was repeated twice.

We then determined the functional potency of the His-ULD/AB complex expressed in the pET-Duet-1 co-expression vector. Since we established that ASXL2-AB stimulates BAP1 deubiquitinase activity by increasing the V_max_, we simply measured the specific activity of BAP1 in the presence of increasing concentrations of His-ULD/AB in order to conserve substrate (Fig. 5C). The His-ULD/AB complex stimulated BAP1 specific activity 4.5 fold using 100 nM Ub-AMC (Fig. 5C). Data plotting the specific activity values against ULD/AB concentration fit well to a one-site dose response curve with a Hill slope of 1.0 and an EC_50_ of 2.8 nM (95%CI: 1.0-7.5 nM) (Fig. 5D), which is within 3-fold of the functional potency we obtained for ASXL2-AB.

### Kinetic studies of the interactions between AB and ULD domains, ULD/AB complex and full-length BAP and BAP-UCH

Using SPR, we tested the affinity and kinetics of BAP1-ULD and ASXL2-AB. ASXL2 was found to bind to GST-ULD, but not GST-UCH or GST alone (Fig. 6A). ASXL2 bound with moderate affinity to GST-ULD, with a steady state K_D_ value of 134 nM (95% CI: 120-149 nM), (Fig. 6 B,C,D). The kinetics of the interaction was relatively fast, with an association rate of 3.8 x 10^4^ M^−1^ s^−1^ and a dissociation rate of 2.4 × 10^−3^ s^−1^ (Fig. 6D). The K_D_ of 67 nM determined by these kinetic parameters were in good agreement with the K_D_ obtained from steady-state analysis.

**Figure 6.**
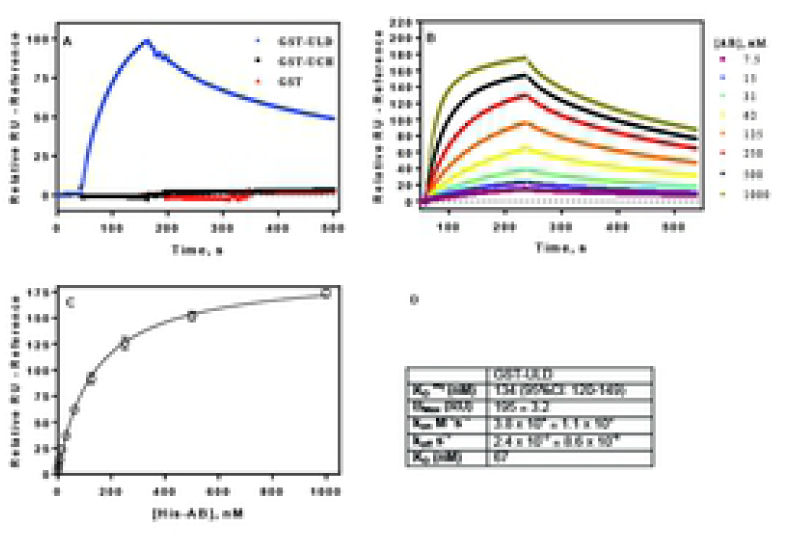
Characterization of the binding of the AB box to the BAP1 ULD domain, as assessed by SPR. **A)** AB box (200 nM) binds to GST-tagged ULD domain of BAP1, but not the UCH domain or GST alone. Data are means of duplicates +/- SEM. **B)** Kinetics of AB binding to GST-ULD. Kinetic parameters were determined from one-site binding model using Biacore evaluation software. Data represent means of duplicate determinations. **C)** Steady-state saturation binding curve of AB binding to ULD. K_D_ and B_max_ values were determined from one-site binding model in GraphPad Prism. Data points are the means +/- SEM of duplicate determinations. **D)** Equilibrium binding and kinetic parameters for interaction of AB and ULD determined from **B** and **C**.

ASXL2-AB by itself did not bind to the UCH domain of BAP1 as determined by SPR (Fig. 6A). Our hypothesis is that both the UCH and ULD domains of BAP1 interact with ASXL2-AB to stabilize the catalytic loop of the UCH domain. Therefore, we investigated the binding of the ULD/AB complex to both the UCH domain and full-length BAP1 using SPR. The ULD/AB complex binds to both full-length BAP1 and GST-UCH with relatively low affinity, but did not bind GST alone (Fig. 7A). The steady-state K_D_ values for full-length BAP1 and GST-UCH were 1910 nM (95% CI: 1600-2400 nM) and 740 nM (95% CI: 580-950), respectively (Fig. 7D). The kinetics of the interaction between the ULD/AB complex and either full-length BAP1 (Fig. 7C) or GST-UCH (Fig. 7B) were characterized by fast association and dissociation rates. The association rates of ULD/AB binding were 3.9 x 10^4^ M^−1^ s^−1^ and 1.9 x 10^4^ M^−1^ s^−1^ for full-length BAP1 and GST-UCH, respectively, and the dissociation rates were 0.033 s^−1^ and 0.044 s^−1^, respectively (Fig. 7B,C,D). The K_D_ values of 850 nM and 2300 nM for BAP1 and GST-UCH, respectively, that were obtained from these kinetic parameters, were in good agreement with those calculated from steady state analysis (Fig. 7B,C,D).

**Figure 7.**
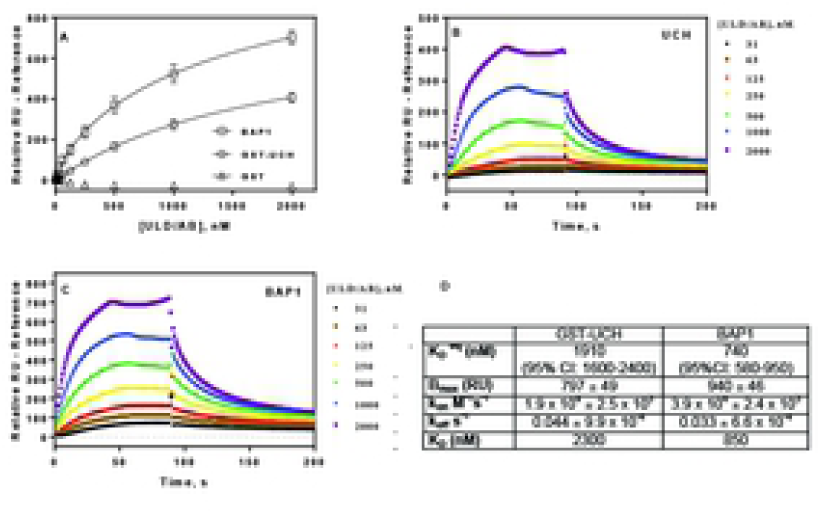
Characterization of the binding of ULD/AB complex to BAP1 and BAP1-UCH domain as assessed by SPR. **A)** ULD/AB complex binds to UCH domain and full length BAP1 but not GST. Steady-state saturation binding curves fit to a one-site binding model. Data are duplicate determinations +/- SEM. **B** and **C**, Kinetics of ULD/AB binding to UCH (**B**) and full length BAP1 (**C**). Kinetic parameters determined from one-site binding model in Biacore evaluation software. Data are means of duplicate determinations. **D)** Equilibrium binding and kinetic parameters for interaction of ULD/AB and UCH or full-length BAP1 determined from **A-C**.

## Discussion

In this report, we have characterized protein-protein interactions between BAP1 and ASXL2 utilizing biochemical and biophysical approaches, as well as enzymatic activity analyses. We have investigated the molecular dynamics, kinetics, and stoichiometry of these intra-molecule and inter-molecule domain-domain interactions. We draw the following conclusions. First, all of the single- or co-expressed and purified recombinant BAP1 and ASXL2 domain/proteins or protein complexes from both bacteria and baculovirus are well-folded in structure and are functionally active. Second, the interaction between BAP1 and ASXL2 is direct, specific, and stable to *in vitro* biochemical and biophysical manipulations. The association of the AB-box greatly stimulates BAP1 deubiquitinase activity. This interaction does not require post-translational modifications. Both bacterial- and baculoviral-expressed BAP1 or BAP1-UCH were enzymatically active and the enzymatic activity increased greatly upon ASXL2-AB box stimulation. A stable ternary complex was formed in UCH/ULD/AB domains. Third, the binding affinity of the ULD domain of BAP to the AB box of ASXL2 is very high with fast association and slow dissociation rates. One molecule of the ULD domain directly interacts with one molecule of the AB Box. Fourth, the formation of this ULD/AB complex with the UCH domain is a single-step event with fast association and slow dissociation rates, indicating that this interaction occurs very rapidly.

To further characterize interactions of domain-domain and tripartite complex between intra-molecule and inter-molecules of BAP1 and ASXL2 proteins, we applied biochemical and biophysical approaches. All these highly purified single- or co-expressed proteins are well structured and capable of folding properly, which allowed us to study the dynamic kinetics of their interactions and stoichiometry of the protein complex association by ITC and SPR (Fig. 2B, Fig. 3A-C). More importantly, the high quality of the bacterial- or baculoviral-expressed proteins and protein complexes are highly functional, which enabled us to perform highly sensitive assays to evaluate deubiquitinase-specific activity of BAP1 and the direct effects of stimulation of ASXL2 on BAP1 enzymatic activity (Figs. 4, 5). These domain-domain interactions and ternary complex interactions were direct and stable (Fig. 3D) and do not require post-translational modifications. We not only were able to reconstitute the tripartite domain complex *in vitro*, but also were able to study the real-time dynamic kinetics of domain-domain and tripartite domain interactions. The binding mode either for AB on ULD, or ULD/AB on UCH are a single-step event with fast association and slow dissociation rates, indicating the interaction is very rapid (Figs. 6, 7). Moreover, the stoichiometry of AB and ULD association occurs via one molecule of AB binding to one molecule of ULD with high affinity (Fig. 3B), which is consistent with the crystal structure of *Drosophila* Calypso / Asx. The stoichiometry of Calypso/Asx was 1:1 molar ratio in low protein concentration, and 2:2 molar ratio in high protein concentration (21). Crystal structure work on the deubiquitinase Calypso, the *Drosophila* counterpart of BAP1, and its activating deubiquitinase adaptor (Deubad) protein partner ASX have provided a structural basis to interpret studies demonstrating that the ASXL1/2 Deubad domains bind tightly to BAP1, and thereby activate the PR-DUB complex by forming a composite binding site for ubiquitin (21). As in our study, Foglizzo et al. (21) showed that mutations at the juncture between DUB, Deubad, and ubiquitin have a deleterious effect on the ability of the PR-DUB to interact with ubiquitin.

We previously showed that the AB box of ASXL2 is the minimal domain required to interact with and stimulate the deubiquitinase activity of BAP1. Mutations in the AB box of ASXL2 or in the ULD domain of BAP1 either partially or completely impacted AB and ULD interaction and UCH ubiquitin hydrolase activity. In this study, we further quantified the AB box protein stimulation on either full-length BAP1 or UCH domain deubiquitinase activity. We observed that ASXL2-AB dose-dependently increased the maximal velocity of BAP1 cleavage of Ub-AMC. Moreover, the ULD/AB complex also increased the maximal velocity of BAP1 cleavage of Ub-AMC in a dose-dependent manner. The data fit well into a one-site dose response equation. The AB box increases the maximal velocity of BAP1-mediated cleavage of Ub-AMC rather than increasing the K_m_ for this substrate. This is consistent with our molecular modeling data suggesting that the AB box does not induce a conformational change in the substrate’s binding pocket, but rather binds to the ULD domain and stabilizes the UCH loop of BAP1. The potency of the AB box for stimulating BAP1 mediated cleavage of Ub-AMC is similar to the concentration of BAP1 in the enzyme assay, which suggests a 1:1 interaction. This is consistent with the ITC results reported here.

Interestingly, the ULD/AB complex, but not the AB box alone, was able to bind the BAP1-UCH domain, as determined by SPR, suggesting that interaction with the ULD domain is essential for stabilizing the UCH domain of BAP1. As the ULD is also found in UCHL5, this makes sense. In addition, most of the affinity for the AB box for BAP1 is through the ULD domain, as this interaction had 10-20 fold higher affinity compared to the affinity of the ULD/AB complex for the UCH domain. These data suggest that the AB box binds the ULD domain first, and this complex then interacts with the BAP1-UCH domain to stimulate enzyme activity.

This is the first quantitative assessment of the inter- and intra-molecular interactions of the BAP1 tumor suppressor and its obligate partner for enzymatic activity, ASXL2, including the mode by which the ASXL2-AB box mediates BAP1 deubiquitinase activity. The tripartite (UCH/ULD/AB) domain-domain interactions described here explain the loss of the BAP1 deubiquitinase activity when tumor-associated mutations in *BAP1* occur outside of the catalytic UCH domain, each failing to productively recruit the AB box to the wild-type BAP1 catalytic site via the ULD, resulting in loss of BAP1 deubiquitinase activity.

In summary, through an integrated use of molecular biology, biochemistry, and biophysics strategies, we have provided evidence to support the molecular mechanism for ASXL2-mediated BAP1 deubiquitinase activity. ASXL functions as a molecular scaffold though its AB box to recruit the ULD domain of BAP1 to transcription factors, which specifically bind to its target genes. Then the UCH catalytic domain of BAP1 ubiquitin hydrolase specifically removes the ubiquitin from histones on chromatin to regulate target genes. ASXL2 not only functions as a molecular scaffold for BAP1 but also greatly stimulates its enzymatic activity. Loss of binding to ASXL2 would dramatically decrease BAP1 deubiquitination activity and thereby lead to BAP1 dependent alterations in chromatin state/gene expression in human cancers and other diseases. Furthermore, small-molecule approaches to reactivate latent wild-type UCH activity of these mutants occurring in a subset of BAP1-mutant cancers might be therapeutically viable.

## The abbreviations used are

UCH: ubiquitin C-terminal hydrolase;
ULD: UCH37-like domain;
PcG: polycomb group (PcG);
(PR-DUB): polycomb repressive deubiquitinase;
ASXH: *Asx* homology domain;
PHD: plant homeo domain;
PRC2: polycomb repressive complex 2;
NLS: nuclear localization signals;
Bac: bacteria;
Bv: baculovirus;
SPR: surface plasmon resonance;
ITC: isothermal titration calorimetry;
CD: circular dichroism;
DLS: dynamic light scattering.

## Acknowledgments

We thank Katherine L.B. Borden and Michael Osborne (University of Montreal) for their advice and suggestions related to this project and manuscript. This work was supported by NCI grants R01CA175691 (J.R. Testa, F.J. Rauscher), P30CA010815 (F.J. Rauscher), P30CA006927 (Fox Chase Cancer Center), R01CA163761 (F.J. Rauscher), and NIH Office of the Director grant K01ES025435 (J.W. Prokop). Support for Shared Resources was provided by P30CA010815 to The Wistar Institute. Also supported by the Jayne Koskinas Ted Giovanis Foundation for Health and Policy (F.J. Rauscher), the Palmira and James Nicolo Family Research Fund (S.B. Malkowicz), Local #14 Mesothelioma Fund of the International Association of Heat and Frost Insulators and Allied Workers (J.R. Testa), Ovarian Cancer Research Fund Alliance (F.J. Rauscher), Samuel Waxman Cancer Research Foundation (F.J. Rauscher), Susan G. Komen grant KG110708 (F.J. Rauscher), and Office of the Assistant Secretary of Defense for Health Affairs, through the Breast Cancer Research Program, under Award numbers W81XWH-17-1-0506, W81XWH-14-1-0235 and W81XWH-11-1-0494 (F.J. Rauscher). This work was also supported by grants 1K22A154600 and 1R03A188439 (E.J. Kennedy).

## Conflicts of interest

The authors declare no potential conflicts of interest.

**Supporting information Figure S1 *BAP1* expression in mouse single cell datasets. A)** The average counts per million of *Bap1* for 20 mouse tissues (x-axis) and the percent of cells in the tissue with an expression greater than 10 counts for *Bap1* (y-axis). **B)** Genes co-expressed in *Bap1* expressing thymus cells. The x-axis shows the log2 fold change of normalized counts for *Bap1* expressing cells and those without *Bap1*. Number of genes identified are shown in the top corners. **C)** Of the *Bap1* expressing cells, the breakdown of those expressing the different *Asxl1-3* genes. The values are for the average among the 20 tissues with the standard deviation shown next to the average. **D)** Breakdown of the *Asxl1* and *Asxl2* values over the 20 tissues from panel **B. E)** Correlation analysis of *Bap1* and *Asxl2* in the 20 tissues. The Spearman’s rank correlation is shown in the legend for each tissue. **F)** Genes that correlate with *Bap1* and *Asxl2* expression in both the liver and pancreas with genes counts shown in the top corners and GO enriched terms colored cyan or red.

## References

1. Jensen DE, Proctor M, Marquis ST, Gardner HP, Ha SI, Chodosh LA, et al. BAP1: a novel ubiquitin hydrolase which binds to the BRCA1 RING finger and enhances BRCA1-mediated cell growth suppression. Oncogene. 1998;16(9):1097–112. Epub 1998/04/07.

2. Nishikawa H, Wu W, Koike A, Kojima R, Gomi H, Fukuda M, et al. BRCA1-associated protein interferes with BRCA1/BARD1 RING heterodimer activity. Cancer Res. 2009;69(1):111–9. Epub 2009/01/02.

3. Misaghi S, Ottosen S, Izrael-Tomasevic A, Arnott D, Lamkanfi M, Lee J, et al. Association of C-terminal ubiquitin hydrolase BRCA1-associated protein 1 with cell cycle regulator host cell factor 1. Mol Cell Biol. 2009;29(8):2181–92. Epub 2009/02/04.

4. Gaytan de Ayala Alonso A, Gutierrez L, Fritsch C, Papp B, Beuchle D, Muller J. A genetic screen identifies novel polycomb group genes in Drosophila. Genetics. 2007;176(4):2099–108. Epub 2007/08/25.

5. Scheuermann JC, de Ayala Alonso AG, Oktaba K, Ly-Hartig N, McGinty RK, Fraterman S, et al. Histone H2A deubiquitinase activity of the Polycomb repressive complex PR-DUB. Nature. 2010;465(7295):243–7. Epub 2010/05/04.

6. Inobe T, Matouschek A. Paradigms of protein degradation by the proteasome. Curr Opin Struct Biol. 2014;24:156–64. Epub 2014/03/19.

7. Isaksson A, Musti AM, Bohmann D. Ubiquitin in signal transduction and cell transformation. Biochim Biophys Acta. 1996;1288(1):F21–9. Epub 1996/08/08.

8. Kadariya Y, Cheung M, Xu J, Pei J, Sementino E, Menges CW, et al. Bap1 Is a Bona Fide Tumor Suppressor: Genetic Evidence from Mouse Models Carrying Heterozygous Germline Bap1 Mutations. Cancer Res. 2016;76(9):2836–44. Epub 2016/02/21.

9. Bott M, Brevet M, Taylor BS, Shimizu S, Ito T, Wang L, et al. The nuclear deubiquitinase BAP1 is commonly inactivated by somatic mutations and 3p21.1 losses in malignant pleural mesothelioma. Nat Genet. 2011;43(7):668–72. Epub 2011/06/07.

10. Harbour JW, Onken MD, Roberson ED, Duan S, Cao L, Worley LA, et al. Frequent mutation of BAP1 in metastasizing uveal melanomas. Science. 2010;330(6009):1410–3. Epub 2010/11/06.

11. Testa JR, Cheung M, Pei J, Below JE, Tan Y, Sementino E, et al. Germline BAP1 mutations predispose to malignant mesothelioma. Nat Genet. 2011;43(10):1022–5. Epub 2011/08/30.

12. Abdel-Rahman MH, Pilarski R, Cebulla CM, Massengill JB, Christopher BN, Boru G, et al. Germline BAP1 mutation predisposes to uveal melanoma, lung adenocarcinoma, meningioma, and other cancers. J Med Genet. 2011;48(12):856–9. Epub 2011/09/24.

13. Carbone M, Yang H, Pass HI, Krausz T, Testa JR, Gaudino G. BAP1 and cancer. Nat Rev Cancer. 2013;13(3):153–9. Epub 2013/04/04.

14. Pilarski R, Rai K, Cebulla C, Abdel-Rahman M. BAP1 Tumor Predisposition Syndrome. In: Adam MP, Ardinger HH, Pagon RA, Wallace SE, Bean LJH, Stephens K, et al., editors. GeneReviews((R)). Seattle (WA)2016.

15. Wiesner T, Obenauf AC, Murali R, Fried I, Griewank KG, Ulz P, et al. Germline mutations in BAP1 predispose to melanocytic tumors. Nat Genet. 2011;43(10):1018–21. Epub 2011/08/30.

16. Peng H, Prokop J, Karar J, Park K, Cao L, Harbour JW, et al. Familial and Somatic BAP1 Mutations Inactivate ASXL1/2-Mediated Allosteric Regulation of BAP1 Deubiquitinase by Targeting Multiple Independent Domains. Cancer Res. 2018;78(5):1200–13. Epub 2017/12/30.

17. Abdel-Wahab O, Adli M, LaFave LM, Gao J, Hricik T, Shih AH, et al. ASXL1 mutations promote myeloid transformation through loss of PRC2-mediated gene repression. Cancer Cell. 2012;22(2):180–93. Epub 2012/08/18.

18. Bainbridge MN, Hu H, Muzny DM, Musante L, Lupski JR, Graham BH, et al. De novo truncating mutations in ASXL3 are associated with a novel clinical phenotype with similarities to Bohring-Opitz syndrome. Genome Med. 2013;5(2):11. Epub 2013/02/07.

19. Hoischen A, van Bon BW, Rodriguez-Santiago B, Gilissen C, Vissers LE, de Vries P, et al. De novo nonsense mutations in ASXL1 cause Bohring-Opitz syndrome. Nat Genet. 2011;43(8):729–31. Epub 2011/06/28.

20. Argote A, Mora-Hernandez O, Milena Aponte L, Barrera-Chaparro DI, Munoz-Ruiz LM, Giraldo-Mordecay L, et al. Cardiovascular Risk Factors and Carotid Intima-Media Thickness in a Colombian Population With Psoriasis. Actas dermo-sifiliograficas. 2017;108(8):738–45. Epub 2017/07/01. Factores de riesgo cardiovascular y grosor de la intima media carotidea en una poblacion colombiana con psoriasis.

21. Foglizzo M, Middleton AJ, Burgess AE, Crowther JM, Dobson RCJ, Murphy JM, et al. A bidentate Polycomb Repressive-Deubiquitinase complex is required for efficient activity on nucleosomes. Nat Commun. 2018;9(1):3932. Epub 2018/09/28.

22. Peng H, Begg GE, Schultz DC, Friedman JR, Jensen DE, Speicher DW, et al. Reconstitution of the KRAB-KAP-1 repressor complex: a model system for defining the molecular anatomy of RING-B box-coiled-coil domain-mediated protein-protein interactions. J Mol Biol. 2000;295(5):1139–62. Epub 2000/02/02.

23. Peng H, Begg GE, Harper SL, Friedman JR, Speicher DW, Rauscher FJ, 3rd. Biochemical analysis of the Kruppel-associated box (KRAB) transcriptional repression domain. J Biol Chem. 2000;275(24):18000–10. Epub 2000/04/05.

